# Optimized Time-segmented Acquisition Expands Peptide and Protein Identification in TIMS-TOF Pro Mass Spectrometry

**DOI:** 10.1101/2024.05.22.591515

**Authors:** Huoming Zhang, Dalila Bensaddek

**Author notes:** **Corresponding Author** Dalila Bensaddek - King Abdullah University of Science and Technology, Corelabs, Thuwal 23500-6900, Kingdom of Saudi Arabia.

## Abstract

We introduce here a novel approach, termed time-segmented acquisition (Seg), to enhance the identification of peptides and proteins in trapped ion mobility spectrometry (TIMS)-Time of flight (TOF) mass spectrometry. Our method exploits the positive correlation between ion mobility values and liquid chromatography (LC) retention time to improve ion separation and resolution. By dividing the LC retention time into multiple segments and applying a segment-specific narrower ion mobility range within the TIMS tunnel, we achieved better separation and higher resolution of ion mobility. This resulted in a substantial increase in peptide identification. In comparison to conventional TIMS methods, which typically scan a static ion mobility range (either from 0.6 to 1.6 or from 0.85 to 1.3), the Seg method significantly enhances the identification of peptides, proteins and average sequence coverage per protein. These findings highlight the potential of the Seg method in expanding the capabilities of TIMS-TOF mass spectrometry, especially for peptide-focused analysis such as post-translational modifications and peptidomics.

## Introduction

In recent years, mass spectrometry (MS)-based proteomics has undergone remarkable advancements, revolutionizing our ability to identify and study proteins within complex biological systems. These developments have resulted in faster, more sensitive, and higher-resolution analyses, providing unprecedented insights into the molecular mechanisms of cells^1^ and their implications for health and disease.^2,3^ However, despite these technological breakthroughs, achieving comprehensive coverage of the proteome, especially for low-abundance proteins, remains a challenge.

Significant progress has been made in improving peptide and protein coverage through advancements in sample preparation and fractionation techniques. Traditionally, proteins were extracted and subjected to either in-gel digestion or in-solution digestion methods. In-gel digestion is simple but labor-intensive and often yields poor peptide recovery, while in-solution digestion commonly involves the use of detergents for efficient protein extraction. However, these detergents can interfere with subsequent enzymatic digestion and mass spectrometric analysis. Modified protocols such as filter-aided sample preparation (FASP),^4^ single-pot, solid-phase-enhanced sample preparation (SP3),^5^ and suspension trap (sTRAP)^6^ have been developed to overcome these challenges. These approaches typically involve the use of strong detergents for protein extraction, followed by the elution of highly pure tryptic peptides. This enables unbiased recovery of proteins, including hydrophobic membrane proteins, and enhances proteome detection sensitivity by removing MS interference. Additionally, selective depletion or targeted enrichment of high-abundance proteins or peptides reduces sample complexity, enhancing the detection of lower abundance proteins. This strategy has been widely employed in the analysis of human body fluids, such as serum, plasma and cerebrospinal fluid, where the removal of the top 20 most abundant proteins enables access to lower abundance proteins.^7^ Moreover, targeted enrichment techniques have proven highly successful in studying post-translational modifications (PTMs). For example, affinity enrichment using Anti-diGly antibody has enabled the identification of over 35,000 ubiquitylation sites from a single data-independent acquisition,^8^ while phosphopeptide enrichment using titanium dioxide (TiO2) and immobilized metal affinity chromatography (IMAC) beads routinely identifies over 10,000 phosphosites.^9,10^ Furthermore, the prefractionation of complex samples into multiple fractions simplifies the analysis and significantly increases peptide and proteome coverage. Remarkably, by combining tissue dissection and extensive HPLC fractionation, draft proteomes covering high percentages of predicted proteins have been reported for organisms such as Human (∼84%)^11^ and Arabidopsis (∼56%^12^ and over 66% in our unpublished data), respectively.

In addition to the optimization of sample preparation, the analysis of these peptide samples has seen remarkable improvement as well. New mass spectrometers, such as high-field orbitraps, offer significantly better resolution and faster speed in acquiring spectral data than their predecessors. Consequently, relatively large proteomes can be quickly and accurately characterized in a single LC-MS injection. By combining the improved MS analyzers with additional ion mobility spectrometry (IMS), further enhancement in both resolution and sensitivity is obtained.^13,14^ IMS separates ions of different structural information based on their size, shape, and charge,^15^ thus providing an additional dimension for separation when integrated into a single mass spectrometry analysis. Trapped ion mobility spectrometry (TIMS) is a relatively new IMS technique that has reversed the concept of classical drift cell analyzers by holding the ions stationary and propelling them with a gas flow through the TIMS tunnel.^16^ It offers several advantages over other IMS techniques such as its ease of integration into MS and offering high ion transmission. In timsTOF Pro MS, Bruker adopted a dual-TIMS analyzer interfacing with Q-TOF MS, where ions are trapped and accumulated in the first TIMS analyzer and subsequently scanned in the second TIMS analyzer before being released into MS. This allows the development of an efficient and highly sensitive workflow called parallel accumulation–serial fragmentation (PASEF)^13^ that synchronizes precursor selection and ion mobility separation and enables sequencing presumably 100% incoming ions.^17^

In a typical PASEF data-dependent acquisition (DDA) experiment, the TIMS analyzer scans ion mobility coefficients (1/*K*0) ranging between 0.6 - 1.6 V s/cm^2^ to cover the nearly entire ion mobility range. This approach can scan up to 600,000 spectra within a 2-hour liquid chromatography (LC) gradient and identify a deep proteome from a complex mixture. Using a narrower ion mobility range, or a combination of multiple mobility windows,^18^ one could increase the depth of the proteomes further. However, narrowing the ion mobility range results in a failure to identify peptides that fall outside the selected range, which poses a disadvantage when studying peptide-centric post-translational modifications. Combining multiple windows significantly lengthens the instrumental analysis time. In this study, based on our previous observation that the ion mobility range has a positive correlation with LC gradient, we developed a time-segmented ddaPASEF acquisition method to analyze complex protein samples. This approach allowed us to identify approximately 15% more proteins and 30% more peptides as compared to the broad-range ion mobility method from commercial HeLa digests. Subsequent application to phosphoproteome analysis enabled us to identify the largest number of phosphorylation sites (>85,000) and phosphopeptides (>150,000) from a HeLa cell culture in a single study.

## Experimental Section

### Standard HeLa Digest

HeLa Protein Digest (Catalog number: 88328, Thermo Scientific) was procured and reconstituted to 200 ng/μL in 0.1% formic acid in HPLC-grade water. Subsequently, this stock solution was further diluted to various concentrations at 100, 50, 25, 10, and 1 ng/μL for analysis.

### HeLa Cell Sample Preparation

Human cervical carcinoma HeLa cells (CCL-2, ATCC) were cultured in DMEM/F-12 media (Gibco, Thermo Scientific, MA, USA) supplemented with 10% fetal calf serum and 1% penicillin/streptomycin antibiotics at 37 °C in a 5% CO_2_ incubator. The cells were cultured in 10-cm petri dishes and sub-culturing was performed when they reached approximately 80% confluence. After reaching 50-60% confluence, the cells were transitioned to serum-free culture medium and allowed to grow overnight. The cells were washed three times with cold PBS and lysed in 500 μl of cell lysis buffer (4 % SDS in 100 mM triethylammonium bicarbonate) supplemented with Halt protease inhibitors and phosphatase inhibitors (Thermo Scientific) and benzonase (5 U/mL, Merck Millipore). The cell lysates were harvested through scraping and transferred to 2-mL Eppendorf tubes. The lysates were subjected to sonication three times for 10 sec each to sheer residual DNA/RNA. The lysates were then centrifuged at 10,000 × *g* for 10 min at 4 °C to eliminate cellular debris. The resulting supernatant containing the proteins of interest was transferred to new Eppendorf tubes, and its protein content was quantified using a microBCA kit (Thermo Scientific).

Approximately 10 mg of proteins from each replicate were purified using methanol/chloroform precipitation and subsequently dried under vacuum. The dried protein pellets were redissolved in an extraction buffer composed of 50 mM triethylammonium bicarbonate and 5% SDS in water, after which proteins were digested using STrap as described^19^. The resulting peptides were desalted using Sep-Pak C18 cartridges (Waters, MA, USA) and dried in a SpeedVac (Thermo Scientific).

### Phosphopeptide Enrichment

Phosphopeptide enrichment was conducted using titanium dioxide microspheres (TiO_2_) following established protocols.^20^ Briefly, the desalted peptides were dissolved in 500 μL of TiO_2_ binding buffer (50% acetonitrile, 2M lactic acid), and incubated for 5 min at room temperature on a Thermomixer shaking at 1,400 rpm. Subsequently, the sample was centrifuged at 10,000 g for 5 min, and the supernatant was transferred to 2 mL Eppendorf tubes containing 500 μL pre-washed TiO_2_ beads slurry by TiO_2_ binding buffer, maintaining a ratio of approximately 5:1 (w/w of TiO_2_ beads to peptides. After a 45-min incubation at room temperature on a Thermomixer with constant shaking at 1,400 rpm, the TiO2 beads containing phosphopeptides were pelleted down by centrifugation at 10,000 g for 5 min. The beads were subsequently washed once with TiO_2_ binding buffer and twice with TiO_2_ washing buffer, composed of 50% acetonitrile and 0.1% trifluoroacetic acid. The TiO_2_ beads were resuspended in 100 μL of TiO_2_ washing buffer and transferred to a C8 packed tip. The sample was centrifuged at 1,000 g for 3 min or until the entire solution passed through. The phosphopeptides were then eluted twice with 100 μL of 5% ammonium hydroxide solution followed by an additional elution using 100 μL of 50% ACN and 2.5% ammonium hydroxide. The combined eluent was acidified with formic acid before drying in a SpeedVac. Finally, the phosphopeptides were desalted using a C18 packed tip.

### Phosphopeptide Fractionation

Phosphopeptide fractionation was performed through high-pH reversed-phase chromatography, as previously described.^21,22^ The peptides were reconstituted in 100 μL of high-pH buffer A (5 mM ammonium hydroxide in water, pH 10) and were subjected to separation on an XBridge Peptide BEH C18 column (Waters) connected to an Accela high-performance liquid chromatography (HPLC) system (Thermo Scientific). A 50-min gradient of buffer B (90% ACN, 5 mM ammonium hydroxide) in buffer A was created for fractionation as follows: start from 0% to 5% B in 3 min, linearly increase to 25% B within 20 min, continuously increase to 40% B in 10 min, ramp up to 70% B in 5 min, hold at 70% for 5 min and followed by 0% B for 8 min. A total of 50 fractions were collected and concatenated orthogonally (mixing various parts of the gradient) into 15 fractions, which were then dried in a SpeedVac.

### LC-MS Analysis

All peptide samples were analyzed using a timsTOF Pro 2 mass spectrometer, coupled with a nanoElute liquid chromatography system (Bruker Daltonik GmbH, Germany). Peptides were directly injected onto a RP-C18 Aurora emitter column (75 µm i.d.× 250 mm, 1.6 μm, 120 Å pore size, aurora gen 2, Ion Opticks, Australia) using a one-column separation method. Three distinct gradient lengths were used for different peptide quantities. Gradient 1, a 30-min gradient, was utilized for the separation of 1-10 ng of peptides at a flow rate of 0.4 µL/min. It began at 2.00 % ACN, rising to 25% ACN in 30 min, followed by ramps to 37% and then 95% B in 2 min each, with a 4-min hold at 95.00 % ACN. Gradient 2, a 60-min gradient, facilitates the separation of 25-50 ng of peptides at a flow rate of 0.3 µL/min. It was initiated at 2.00 % ACN, with a linear increase to 25% ACN over 60 min, followed by a climb to 37% ACN and finally 95% ACN in 4 min each, with an 8-min hold at 95.00 % B. Gradient 3 is a 90-min gradient with a flow rate of 0.25 µL/min, which was used for peptide quantities ranging from 100 to 200 ng. The gradient began at 2.00 % ACN and progressed to 25% ACN in 90 min, followed by a climb to 37% ACN and finally 95% ACN in 4 min each, with a 7-min hold at 95.00 % ACN. The column temperature was set to 50 °C. Peptides were introduced into the mass spectrometer via a CaptiveSpray nano-electrospray ion source (Bruker Daltonik GmbH) at an electrospray voltage of 1.6 kV, an ion source temperature of 180 °C and a dry gas of 3 l/min.

Data collection employed PASEF scans as previously described,^13^ with minor adjustments. Briefly, a total of 9 PASEF ramps targeting precursors with a charge between 0-5 and a cycle time of 1.26 s. Each PASEF ramp was set to a ramp time of 120 ms. Three distinct TIMS scan ranges were evaluated for their ability to identify peptides and proteins. Method 1 employed a narrow range of 0.85 to 1.30 Vs cm^−2^ (1/K_0_), Method 2 used a conventional range of 0.60 to 1.60 Vs cm^−2^ (1/K_0_), and Method 3 (termed as “Seg”) employed 10 various ranges. The collisional energy increased linearly from 20.0 eV at 0.60 (1/K_0_) to 59.00 eV at 1.60 Vs cm^−2^ (1/K_0_). Both MS and MS/MS spectra were recorded within a scan range of 100 - 1700 *m/z*. TIMS accumulation time was set to 100 ms, with a target precursor intensity was 15,000 arbitrary units (au) and a minimum threshold of 1500 au. Active exclusion was set to 0.4 min. Absolute intensity thresholds were established at 10 for mass spectra peak detection and 5000 for mobilogram peak detection.

### MS Data Processing

MS raw files (.d) were searched against a concatenated target-decoy protein database. The database included the Uniprot human database (20,385 entries, updated on 20 November 2022), common 48 contaminants, and their reversed sequences. HeLa searches were performed using MSFragger (v3.7)^23-25^ via FragPipe (v19.0). Search parameters included cysteine carbamidomethylation as a fixed modification and methionine oxidation, N-terminal protein acetylation, and deamidation at asparagine and glutamine residues as variable modifications. The enzyme limits were set at trypsin cleavage from at least one end of the peptide (termed “semi-tryptic” search) or both ends (“tryptic”), allowing for a maximum of two missed cleavages. Positive peptide identification was required to contain a minimum of seven amino acids. Precursor ion and fragment ions mass tolerances were set at 15 ppm and 0.05 Da, respectively. The false discovery rates (FDRs) of peptide-spectral match (PSM) and protein identification were all set to 0.01. Label-free quantitation was carried out using IonQuant (v1.8.10),^26^ which was integrated into FragPipe.

Similarly, for phosphorylation site analysis, both tryptic and semi-tryptic searches were performed using MSFragger due to its rapid processing speed. In addition to the previously mentioned parameters, variable modifications included phosphorylation at serine, threonine, and tyrosine residues. PTMProphet^27^ was used to assess site localization, with a site probability threshold of 0.75 established for confident phosphosite assignment.

## Results

### Temporal Ion Mobility Profiling over LC elution

In addition to advancements in sample preparation and chromatography separation, effective ion traps and separation in MS play a pivotal role in the comprehensive sequencing of complex peptide mixtures. The integration of ion mobility separation with mass spectrometry substantially enhances peak capacity and deconvolutes complex masses,^28^ thereby facilitating proteome-wide analysis of protein alterations within samples in a single-shot LC-MS run.

In our efforts to optimize ion separation within the timsTOF MS, we first conducted an examination of peptide elution patterns across a spectrum of sample amounts, using three commonly used LC gradients (30, 60 and 90 mins), and the routine PASEF acquisition method. As shown in Figure 1, all peptides eluted out within the 30 min gradient at 5 ug injection whereas with 10 ug, a small number of peptides accumulated and eluted only at high organic buffer (95% B), suggesting that the 30-min gradient yields optimal results for samples containing 10 ng or less. Similarly, the 60- and 90-min gradients offered effective peptide separation for sample amounts up to 50 ng and 200 ng respectively. It is important to note that shorter LC gradients, while being efficient, tend to have lower peak capacities and are susceptible to co-elution of numerous peptides, which can lead to ion suppression during the ionization process.^29^ Hence, it is recommended to select a gradient length that corresponds to the specific sample amount to maximize identification.

**Figure 1.**
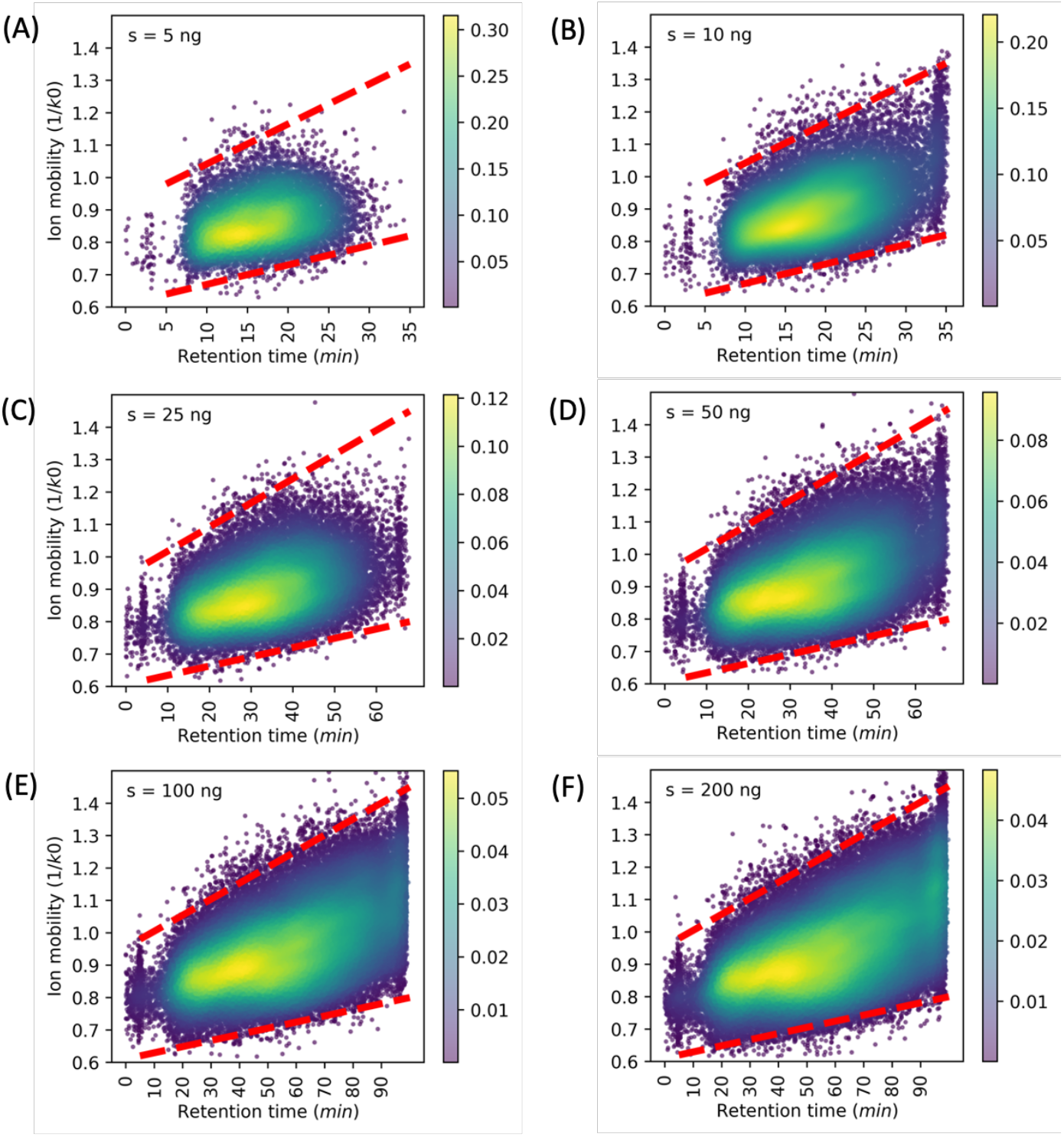
Temporal peptide ion mobility profiles over the 30-, 60- and 90-min LC-gradients. Peptide samples were analyzed using conventional PASEF method with wide range of ion mobility windows (0.60 to 1.60 Vs cm^−2^).

Interestingly, we found a positive correlation between elution time and ion mobility (1/k0 value) of identified peptides, with ion mobilities increasing over elution time, across all tested conditions (Figure 1). Similar observations have been reported earlier that emphasized the significance of elution time as one of critical parameters for accurate prediction of ion mobility.^30^ Moreover, we found that peptides eluted at any retention time fall within a narrow ion mobility window. For example, the majority of peptides have an ion mobility range of 0.63-1.05 at 20 min retention time of the 90 min LC gradient). This time-bound narrow ion mobility property has implications for enhancing both the precision and resolution of peptide identification in complex samples when compared to the common approach of employing a broad TIMS scan range (0.6-1.6, 1/k0) across the entire gradient.

Building on this, we designed a time-segmented acquisition method (termed “Seg”). In this novel approach, ions that initially accumulated in the first TIMS are scanned in the second TIMS using multiple time-dependent narrower ion mobility segments but covering nearly all ions before being released into the mass analyzer. For instance, a total of 9 segments were used for a 30-min gradient separation, with each having a narrow TIMS scan range of ion mobility, such as 0.66-1.02 for the first 5 min of data acquisition, followed by 0.68-1.04 for the next 5 min, and ultimately 0.80-1.35 for the last 5 min of acquisition, as depicted in Figure 2A. Similar approaches are applied to experiments with longer LC gradients (Figure 2B, C). This strategy is anticipated to improve the ion mobility separation and increase the TIMS capacity for ion clusters destined for MS analysis, potentially resulting in a greater number of peptide and protein identifications.

**Figure 2.**
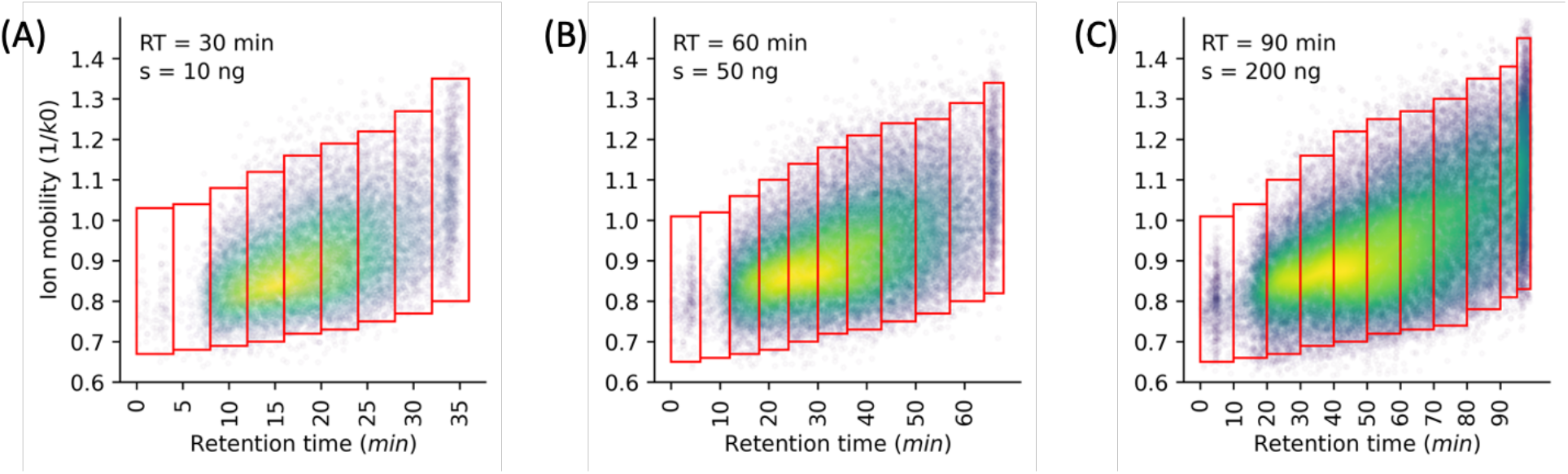
Design of the time-segmented methods for three LC-gradients (A: 30 min, B: 60 min and C: 90 min) according to their ion elution profiles. A total of 9-11 segments with each having a narrower ion mobility scan window as compared to the commonly used range of 0.6-1.6 1/k0.

### Time-segmented Acquisition Increases Identification

To verify our hypothesis, we compared this time-segmented acquisition (Seg) to two commonly used methods with TIMS scan ranges of 0.85-1.3 (referred to as “Narrow”) and 0.6-1.6 Vs cm^−2^ (1/K0, referred to as “Wide”) under different gradient lengths and varying sample amounts. As expected, an increased number of proteins was identified from higher sample injection amounts across all methods, with approximately 2,230 to 7,200 proteins identified from Hela digest loads between 5 ng and 200 ng (Figure 3A). The Seg approach consistently identified the highest number of proteins across all sample loads, outperforming the Wide method by consistently identifying 10-15% more proteins. It also identified a higher number of proteins than the Narrow method albeit in small increments. The superiority of the Seg method became more pronounced when analyzing peptide identification. As shown in Figure 3B, the Seg approach identified about 20-30% more peptides than the Wide method, and about 25-70% more than the Narrow method across all conditions. This demonstrates that the Seg method could increase not only the depth of protein identification but also the sequence coverage of identified proteins. It is worth noting that although the Narrow approach recovered fewer peptides than the Wide method, it detected a greater number of proteins. This discrepancy can be attributed to the Narrow approach’s superior sensitivity for identifying lower abundance peptides, resulting from the narrower scan range and reduced ion complexity.

**Figure 3.**
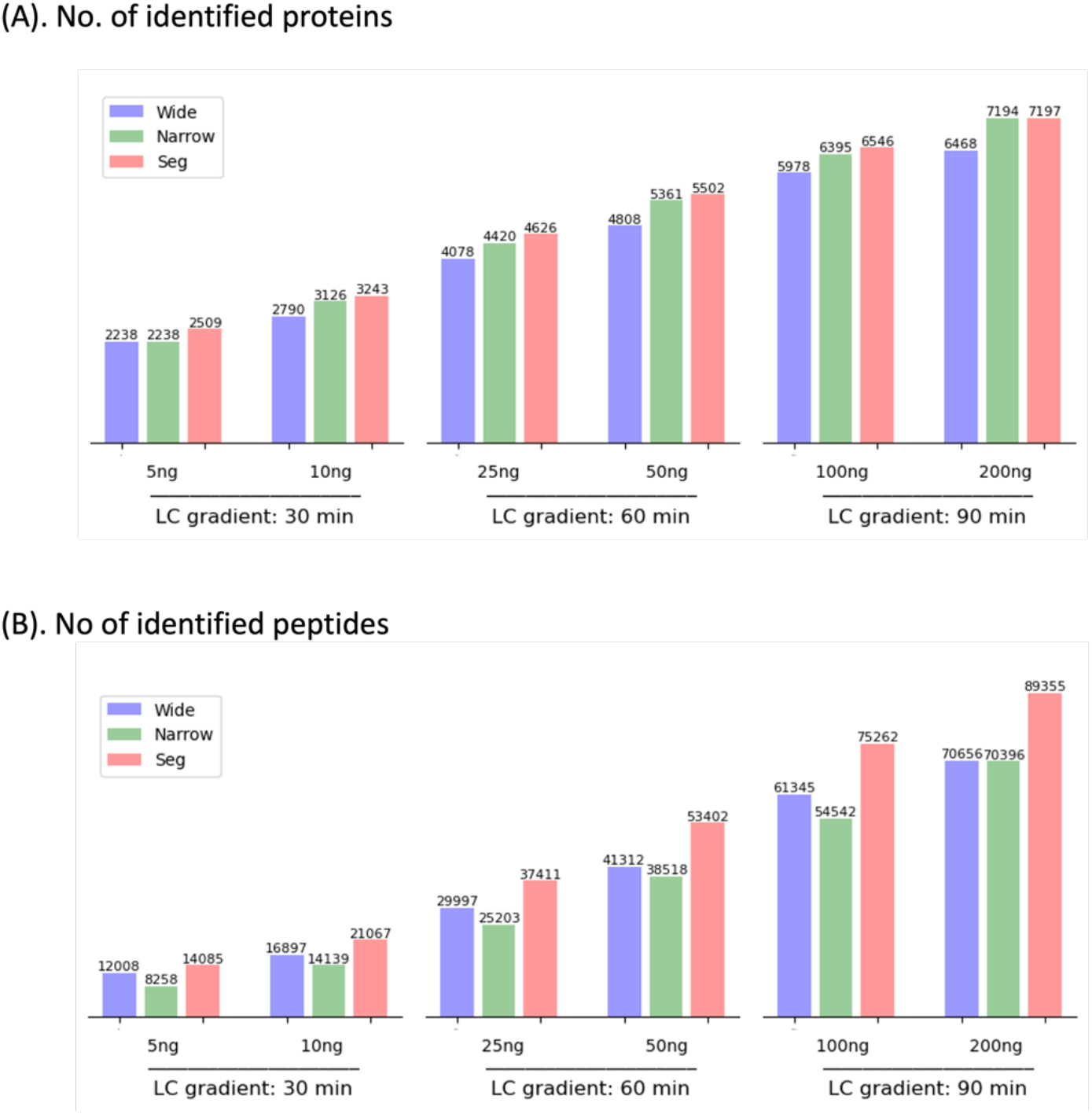
The numbers of protein (A) and peptide identifications (B) from 5 - 200 ng of hela digest using the three Tims scan acquisition methods: the Wide, Narrow and Seg method.

Next, we examined protein and peptide identification from 200 ng of Hela digest across the three methods. Of the total 7,694 proteins identified, approximately 80% of proteins were consistently detected across all methods, indicating a high degree of overlap in protein identification (Figure 4A). Both the Seg and Narrow methods identified over 300 unique identifications whereas the Wide method identified fewer than 100. The identification overlap between Seg and Narrow methods is approximately 90%, suggesting a similar performance between these two methods, while the overlap between Wide and Seg/Narrow is slightly lower at around 85%. Subsequently, we created line plots to visualize the logarithmic intensity versus protein abundance rank. It was observed that the Seg and Narrow methods exhibited comparable protein detection across different abundances (Figure 4C) and demonstrated higher sensitivity in detecting more proteins at relatively low abundance compared to the Wide method (Figure 4E). This suggests that the Seg and Narrow methods are more effective in identifying proteins present in lower quantities.

**Figure 4.**
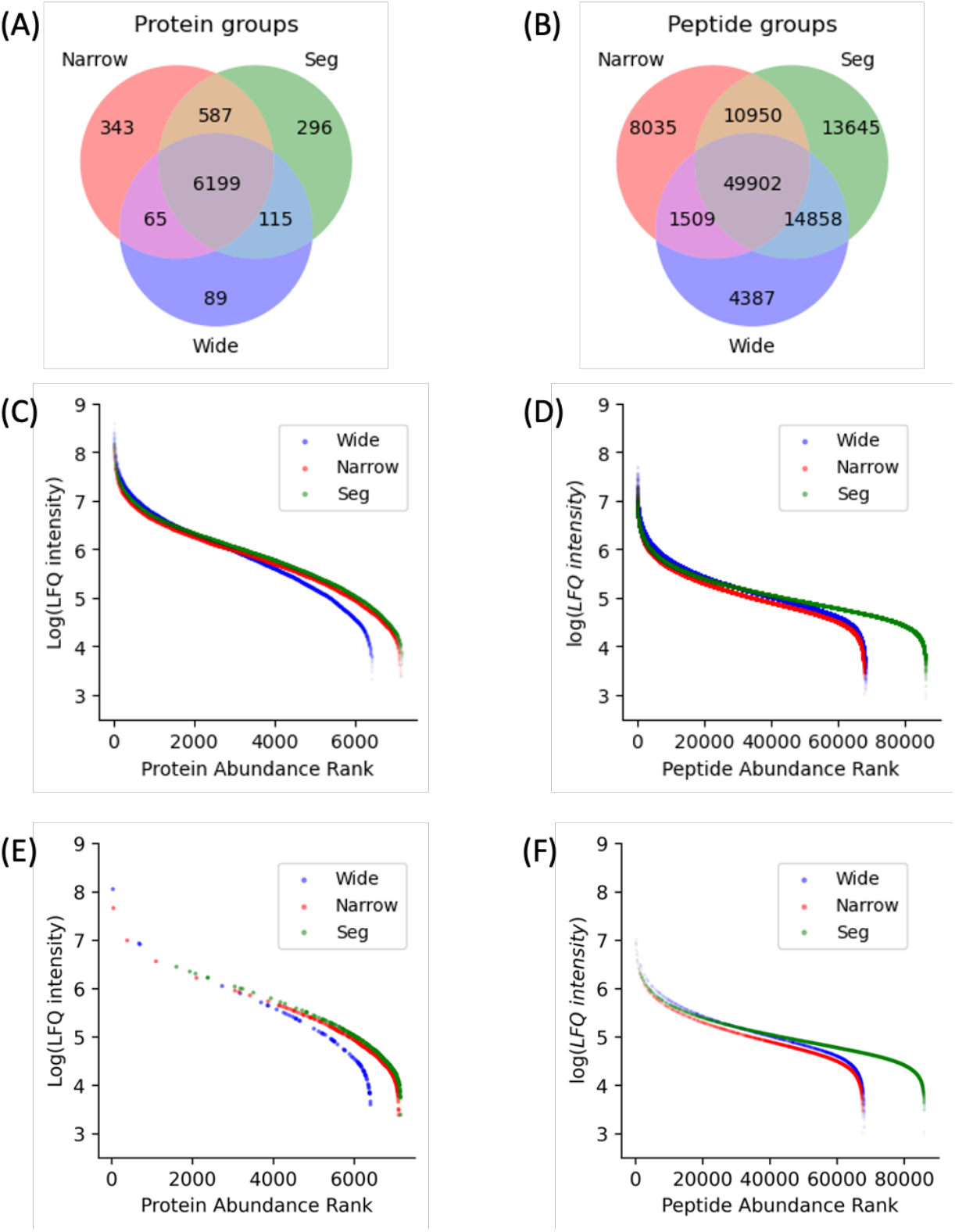
Comparison of protein and peptide identifications from 200 ng of HeLa digest among the three MS-acquisition methods. (A) Venn diagram shows the unique and overlap protein identifications. (B) Venn diagram of peptide identifications. (C) Protein detection across abundances. (D) Peptide detection across abundance. And abundance of the unique detection of proteins (E) and peptides (F), the logarithmic intensity of detected proteins and peptides plotted against their rank by abundance.

In our peptide analysis, a total of 103,286 peptides were identified, with only 48.3% of these peptides being common among all three methods (Figure 4B). The Seg method, which yielded the highest number of unique peptides (13,645), contributed significantly more unique identifications than the combined total from the Wide and Narrow methods. Additionally, the Seg method identified a greater number of low abundance peptides, further highlighting its sensitivity to capture peptides present at lower levels. While the Wide and Narrow methods shared a similar abundance distribution, the Narrow method recovered peptides with slightly lower intensities compared to the Wide method (Figure 4D). Furthermore, like unique proteins, the identified unique peptides were mostly at lower abundance levels (Figure 4F). This suggests that the unique peptides identified by the different methods are predominantly present in smaller quantities. Together, our findings demonstrate that the Seg method outperforms the other two methods in both protein and peptide identification. It exhibits superior performance in detecting proteins across different abundance ranges and shows higher sensitivity to proteins and peptides at lower abundance levels. The Seg method offers exceptional efficiency in peptide identification, making it the ideal choice for comprehensive analysis, especially in samples with low-abundance proteins and peptides.

### Time-segmented Acquisition Enable Accurate Quantitation

To evaluate the quantitative performance of the workflow, we analyzed the results of 200 ng HeLa runs in triplicate using a label-free quantitative approach. In the Wide method, approximately 6,030 proteins were common between any two runs, whereas over 6,740 overlap proteins were identified between any two replicates for the Seg and Narrow methods. These triplicate analyses from each method demonstrated excellent reproducibility, with pairwise Pearson correlation coefficients (R) between replicates of greater than 0.97 over 4.5 orders of magnitude in protein abundance, indicating accurate quantification and that any observed differences can be reliably attributed to actual sample variance. Furthmore, the median coefficient of variation (CV) of protein group quantities showed similarly low values of <1% in both the Wide (Figure 5C) and Seg (Figure 5D) methods, indicating comparable high quantitation accuracy of the Seg method to that of the Wide method.

**Figure 5.**
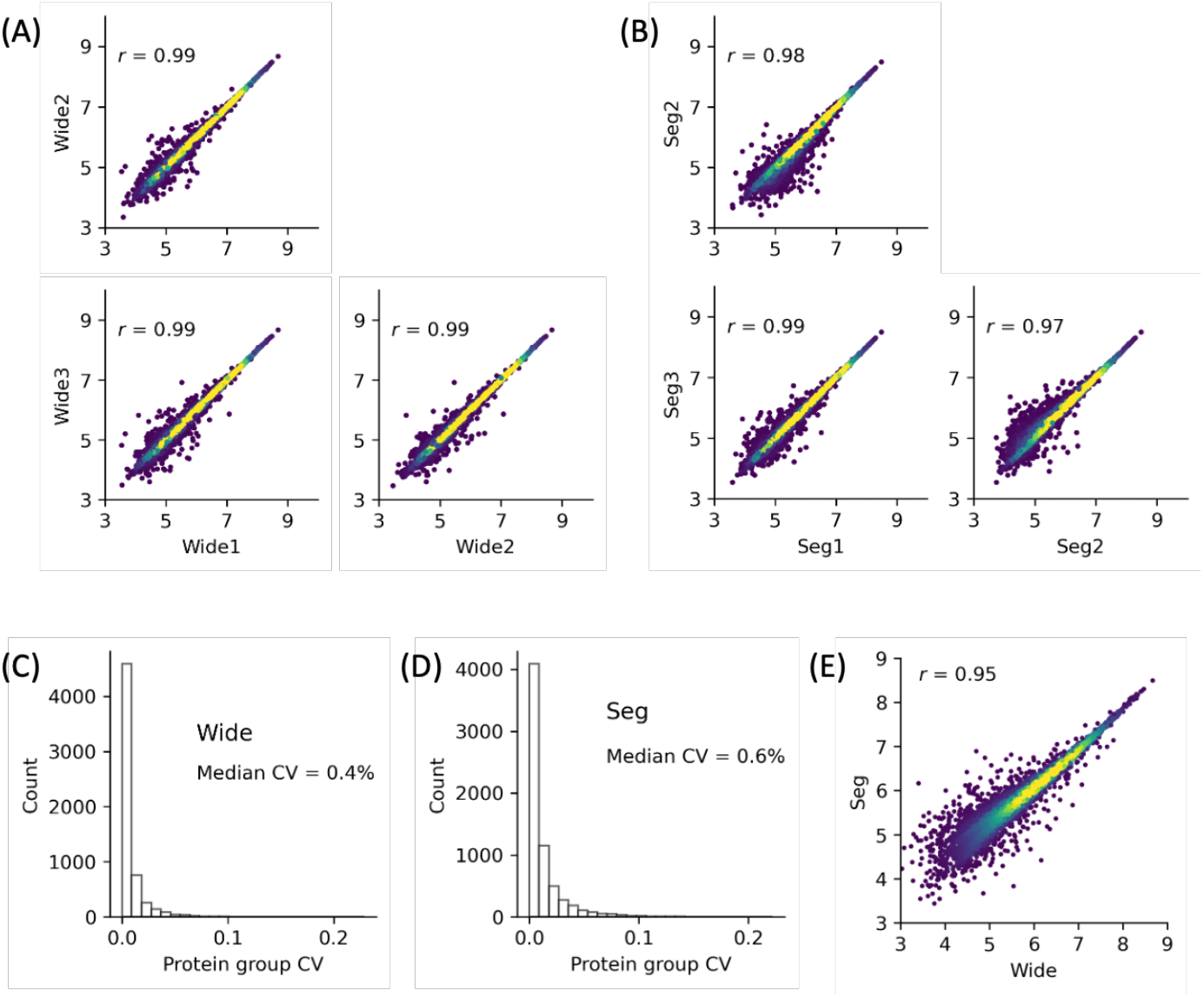
Label-free quantification of protein identifications. The quantitative correlation (protein intensities) between any two replicates in the Wide (A) and Seg methods (B). The coefficients of variation (CVs) for protein quantities in three replicates for the Wide (C) and Seg method (D). (E) Protein quantitative correlation of proteins identified by the Wide and Seg method. The Pearson correlation coefficients are indicated.

We also performed a direct comparison between Seg and Wide methods for non-normalized protein intensities. The Pearson correlation coefficient of common identifications (n=6,224) was high, at over 0.95 (Figure 5E), illustrating the overall similarity in quantitation achieved by the two methods. Together, we demonstrate that the Seg method was able to increase the number of identifications while maintaining high quantitation accuracy.

### Phosphoproteome Analysis

Since the Seg method substantially outperforms other methods in peptide identification, we anticipate it as an ideal choice for studying all types of peptide-centric analysis, such as phosphorylation site analysis. To test this hypothesis, we compared the Wide and Seg method for an in-depth phosphoproteome analysis. A total of 15 fractions from high pH (HpH) chromatography of TiO2 enriched peptide samples were analyzed using the two TIMS scan methods. As expected, the Seg method consistently identified over 10% more phosphosites across all fractions compared to the Wide method, underscoring its enhanced detection capabilities (Figure 6A). While both methods shared substantial cores (131,526 peptides and 50,204 phosphosites, respectively), the Seg method significantly outperformed the Wide method by identifying nearly twice the number of unique peptides (79,153 vs. 37,929) (Figure 6B) and and unique phosphosites (21,263 vs. 11,118) (Figure 6C), highlighting its sensitivity and ability to detect a wider range of peptide modifications. Together, we were able to present the largest number of phosphosite identifications (82,585) from non-stimulated HeLa cell culture in a single study (Table S1). Furthermore, by using a semitryptic search strategy, an additional 4,584 phosphosites were identified from semitryptic peptides, bringing the total number of phosphosites identified in this study to 87,169 sites (Figure 6D, Table S2). To assess the label-free quantitation accuracy, we plotted the log-transformed intensities of phospho-peptides and -proteins identified by both methods. High correlation coefficients (r >= 0.94) were found for both phosphopeptides (Figure 6E) and phosphoproteins (Figure 6F), indicating that increased peptide recovery in the Seg method did not compromise precision in quantitation.

**Figure 6.**
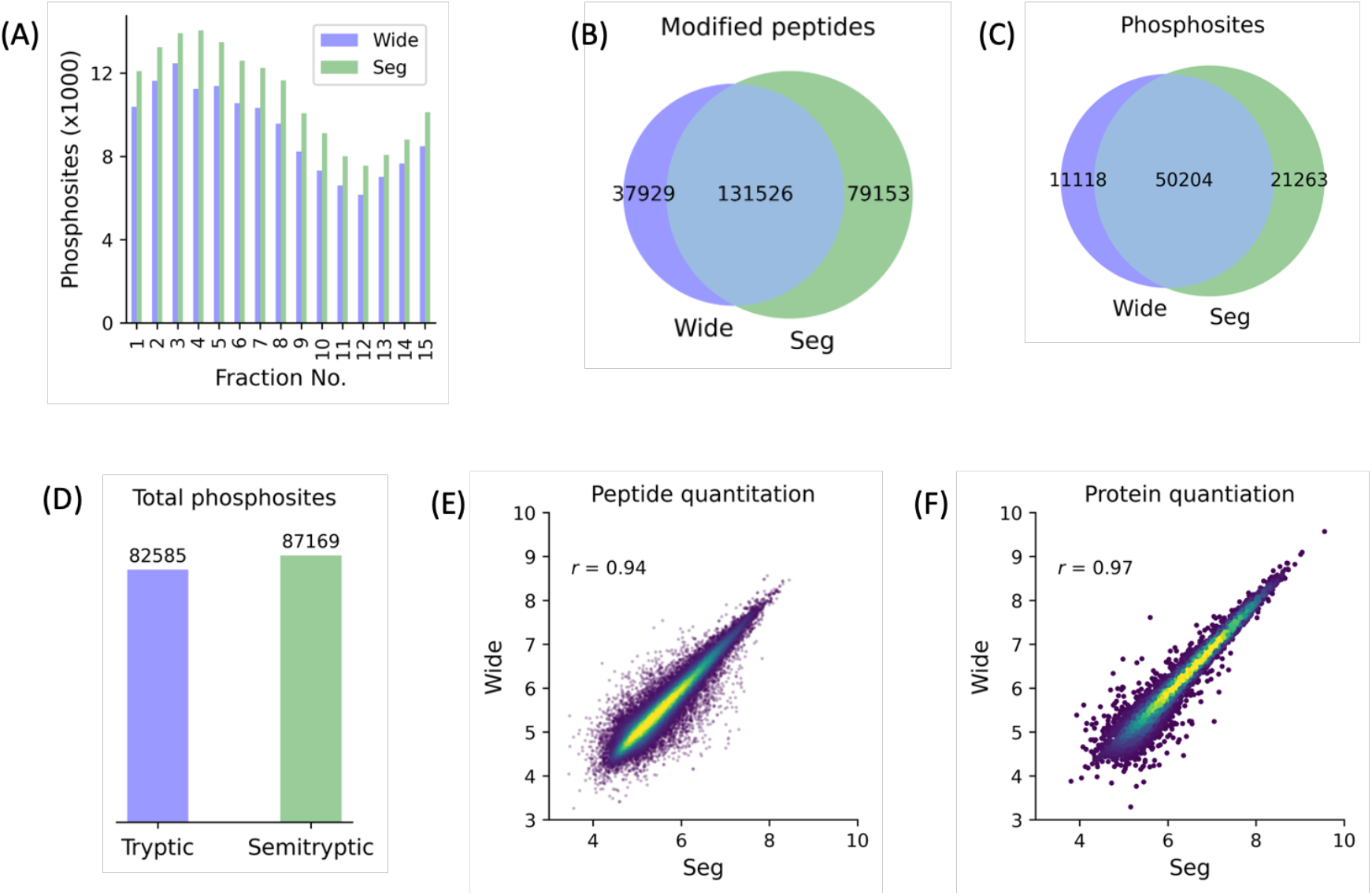
Comparison of the Wide and Seg methods for the phosphoproteome analysis of cultured HeLa cells. (A) Distribution of identified phosphosites across HpH fractions, demonstrating enhanced phosphosite identification by the Seg method. (B) Comparative analysis of modified peptides identified by each method, with the Seg method identifying twice as many unique peptides as much as the Wide method. (C) Venn diagram illustrating the overlap and unique identification of phosphosites. (D) Total number of identified phosphosites from tryptic and semi-tryptic database searches respectively. (E, F) Quantitative comparison of log-transformed intensities of identified phosphopeptides (E) and phosphoproteins (F) between the Wide and the Seg, demonstrating a high correlation coefficient (r) and strong agreement in quantification between the two methods.

In summary, the Seg method exhibits a clear advantage over the Wide approach for phosphoproteomic analysis by reliably identifying more phosphosites and modified peptides, including those of low abundance and non-tryptic origin, thereby providing a more comprehensive view of the phosphoproteome. This confirms the compelling advantages of the Seg acquisition method over the Wide approach for analyzing peptides with and without post-translational modifications.

### Narrow Ion mobility Window Increases TIMS Resolution

To explore the underlying difference among TIMS methods, we generated Extracted Ion Chromograms (XIC) for three “house-keeping” ions of Hela digest (m/z values: 599.76, 566.76 and 663.85) (Figure 7A). These ions, with varying abundances, are consistently detectable and evenly distributed across LC gradients, making them ideal targets for routine quality checks of peptide retention time. We then compared the TIMS resolutions of these representative XIC (Figure 7B). Clearly, both Narrow and Seg methods demonstrated a minimum of 20% higher TIMS resolution compared to the Wide approach (with p<0.05) for all three XIC. This improvement suggests that narrowing the TIMS scan, while maintaining the same electric voltage ramping time, enhances resolving power. Interestingly, the Narrow and Seg displayed similar TIMS resolving power, with one peptide exhibiting slightly higher resolution in the Seg method and the other two showing marginally better resolution in the Narrow method. These comparisons highlight the potential benefits of fine tuning TIMS scan windows to optimize the performance of proteomics experiments.

**Figure 7.**
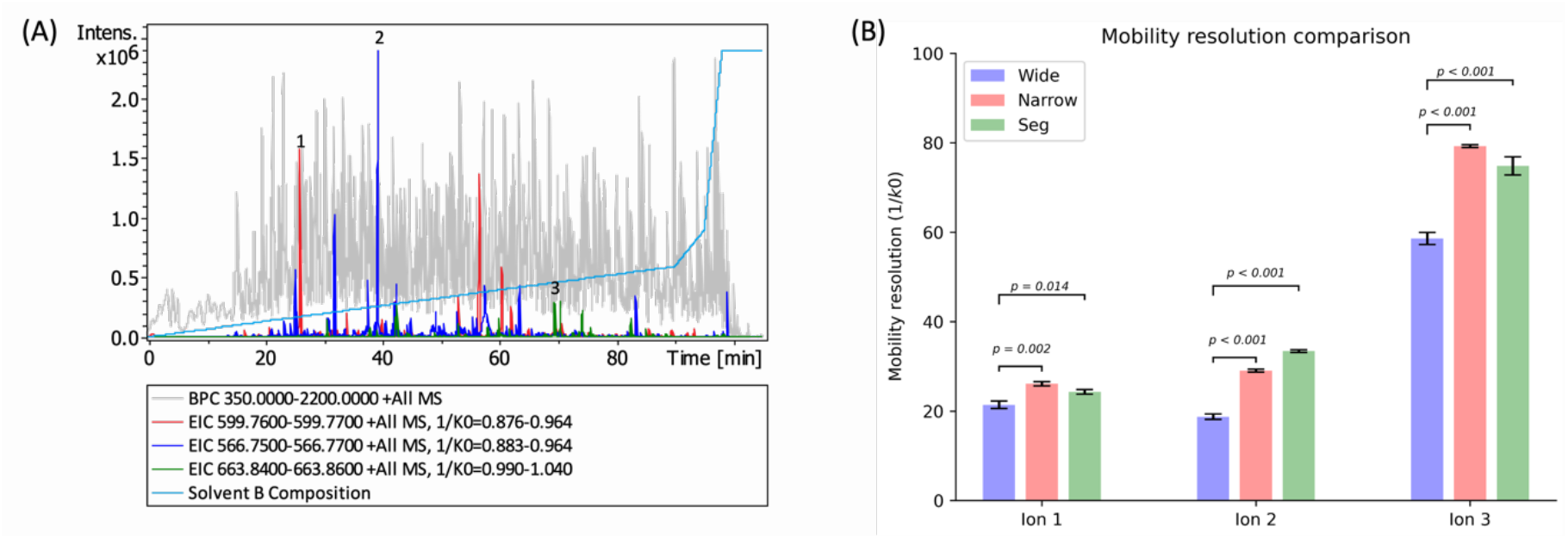
TIMS resolution of three representative ions. (A) Representative LC-MS spectra indicating base peak chromogram, Extracted Ion Chromograms (XIC) for three ions, and LC running gradient (solvent B composition). (B) Comparative bar graph showing the differences in TIMS resolution of three XIC from the Wide, Narrow and Seg methods.

## Discussion

Ion separation prior to MS analysis is crucial for in-depth proteome analysis. Ion mobility MS allows an additional dimension to ion separation, leading to more thorough coverage in high throughput proteomics analysis. With the TimsTOF pro instrument, the ion mobility tunnel is about 100 mm long and can be electrically divided into two equal sections: the ion trap region and the TIMS scan segment, facilitating efficient on-line PASEF analysis.^13^ When ions enter the TIMS tunnel, they undergo longitudinal separation, with ions of lower mobilities positioned closer to the tunnel’s exit and ions of higher mobilities concentrated near the entrance, before being sequentially released to MS analysis.^31^ The effectiveness of ion capacity and separation in the TIMS tunnel depends heavily on the alignment between the tunnel’s length and the range of ion mobilities, considering the impact of space charge effects on ions.

Theorectically, the use of a TIMS scan range of 0.6-1.6 (ion mobility) can be ideal for scanning a cluster of ions having similar mobility range. However, this range may not be suitable for peptide ion clusters with significantly smaller mobility ranges, such as between 0.8 and 1.3, which is the case in actual analysis of complex peptide samples (Figure 1). Although the ion mobilities range from 0.6 to 1.5 across the whole LC gradient, the fragmented and identified ions concentrate on much narrower range at any small elution time segment (Figure S1). Use of the Wide approach reduces TIMS separation efficiency as a large proportion of the scan time is used to acquire spectra on ion mobility regions with no ion of interest present. In this study, we introduced an improved TIMS scan approach with time-segmented acquistion (Seg) for enhancing TIMS separation and resolution of complex peptide mixtures. This approach utilized the temporal property of peptide ion mobility associated with LC elution time. Each TIMS segment scans a smaller ion mobility window based on the specific mobility characteristics of the peptide ion clusters. The mobility windows are nearly 50% smaller than the Wide strategy, resulting in increased TIMS length space for the same cluster of peptide ions, thereby reducing space charge effects and improving ion separation and resolution. Moreover, this approach reduces non-peptide ions or unidentifiable ions during TIMS scans, thus increasing MS time on fragmentation of identifiable peptides. Our Seg approach further enhances the ion mobility resolution by lowering the voltage ramp speed (equivalent to narrowing the mobility range while maintaining the same elution ramp time). The improvement leads to enhanced sensitivity and specificity, particularly beneficial to low abundance peptide ions within ion clusters at a wide dynamic range. Indeed, we typically observed over 20% more peptide identifications for analyzing varying loads of Hela digest (Figure 3B). Notably, the majority of additional peptides and proteins are in low abundance (Figure 4F). In comparison to the Narrow approach with a fixed ion mobility range (0.85-1.3), the Seg method uses a similarly narrow mobility window but scans nearly all peptide ions, demonstrating superiority for peptide-centric analysis with significantly deeper coverage. Furthermore, the Seg method achieves comparable coverage without the need for extended instrumental measurement times typically required when using the mobility window fractionation approach.^18^

In conclusion, the Seg method is a valuable approach for enhancing ion separation and resolution in complex peptide mixtures analyzed using TIMS. By employing narrowed ion mobility windows, the Seg method increases resolving power, reduces ion suppression, enhances sensitivity, and improves data quality. This method is particularly suitable for peptide-focused analyses, such as post-translational modifications and immunopeptidomics, as well as analyses requiring de novo sequencing.

## Associated Content

The Supporting Information, MS raw data and their search results are available free of charge at ProteomeXchange Consortium via the PRIDE^32^ partner repository with the dataset identifier PXD051790.

## Author Information

## Author Contributions

The manuscript was written through contributions of all authors. / All authors have given approval to the final version of the manuscript.

## Notes

The authors declare no competing financial interest.

## Acknowledgment

We thank the director of the Analytical Chemistry Core Laboratory, Dr Maan Amad, for his support in this project.

---

**Figure S1.**
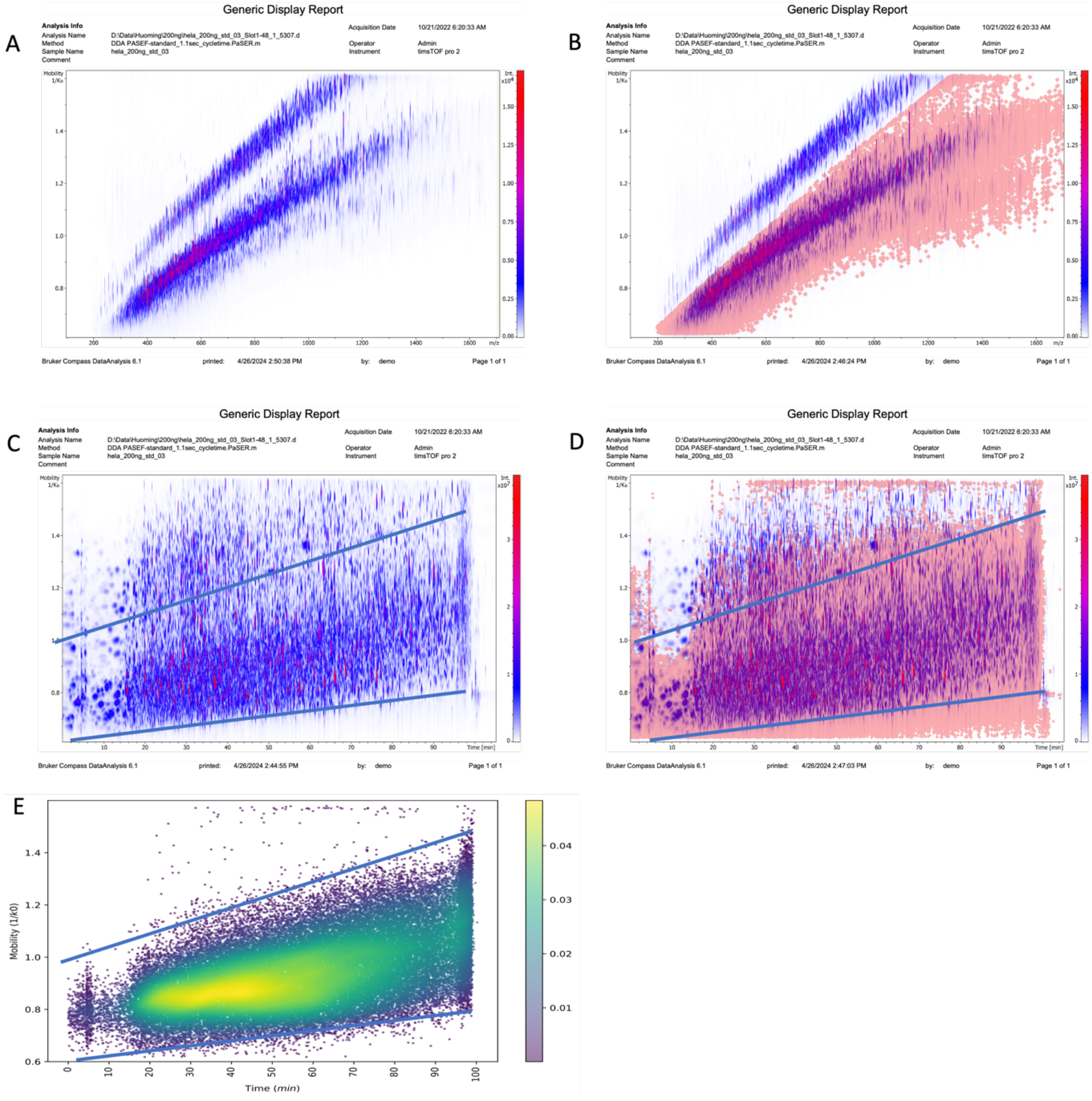
Heatmap of ion elution profiles for a typical 200 ng Hela sample with 90-min gradient. (A) Ion mobility (1/k0) vs ion mass-to-charge (m/z) for all ions. (B) Ion mobility (1/k0) vs ion mass-to-charge (m/z) for all ions, with MS/MS of ions highlighted in a light-red square. (C) Ion mobility (1/k0) vs LC retention time for all ions. (D) Ion mobility (1/k0) vs LC retention time for all ions, with MS/MS of ions highlighted in a light-red square. (E) Ion mobility (1/k0) vs LC retention time for the ions identified as peptides via database search. Clearly, most of ions outside the two lines are either singly charged or non-identifiable ions (C-E).

## Notes

### Competing Interest Statement

The authors have declared no competing interest.

